## Supplemental Figure S1 for "Effective population size does not explain long-term variation in genome size and transposable element content in animals"

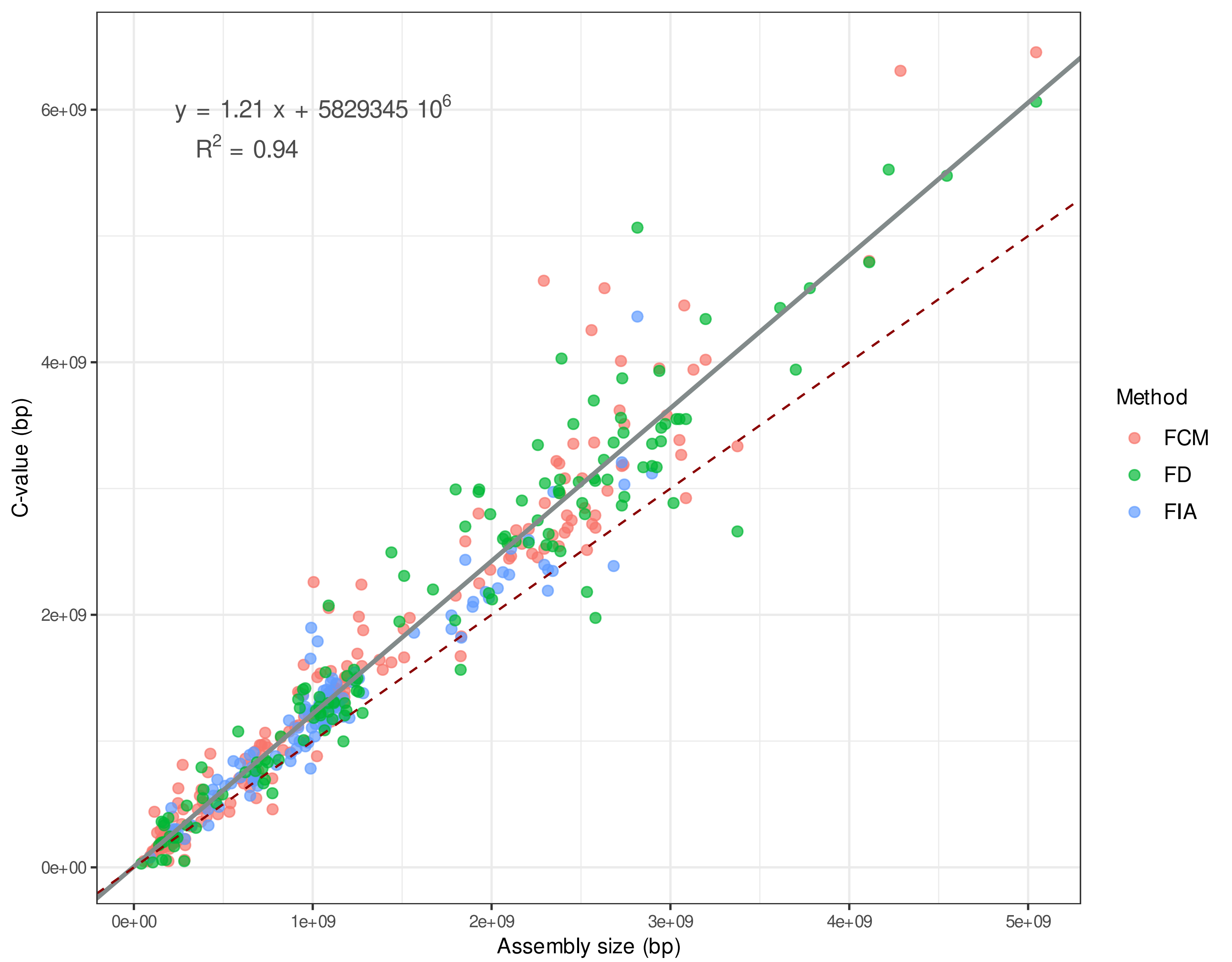


Supplemental Figure S1. Correlation between assembly sizes and C-values for 365 species with contig N50 ≥ 50 kb. The grey slope corresponds to the WLS used to predict the expected C-values (reported in the equation). The dark-red dashed slope marks the hypothetical 1:1 relationship. FCM = Flow Cytometry, FD = Feulgen Densitometry, FIA = Feulgen Image Analysis.
