## Supplemental Figure S2 for "Effective population size does not explain long-term variation in genome size and transposable element content in animals"

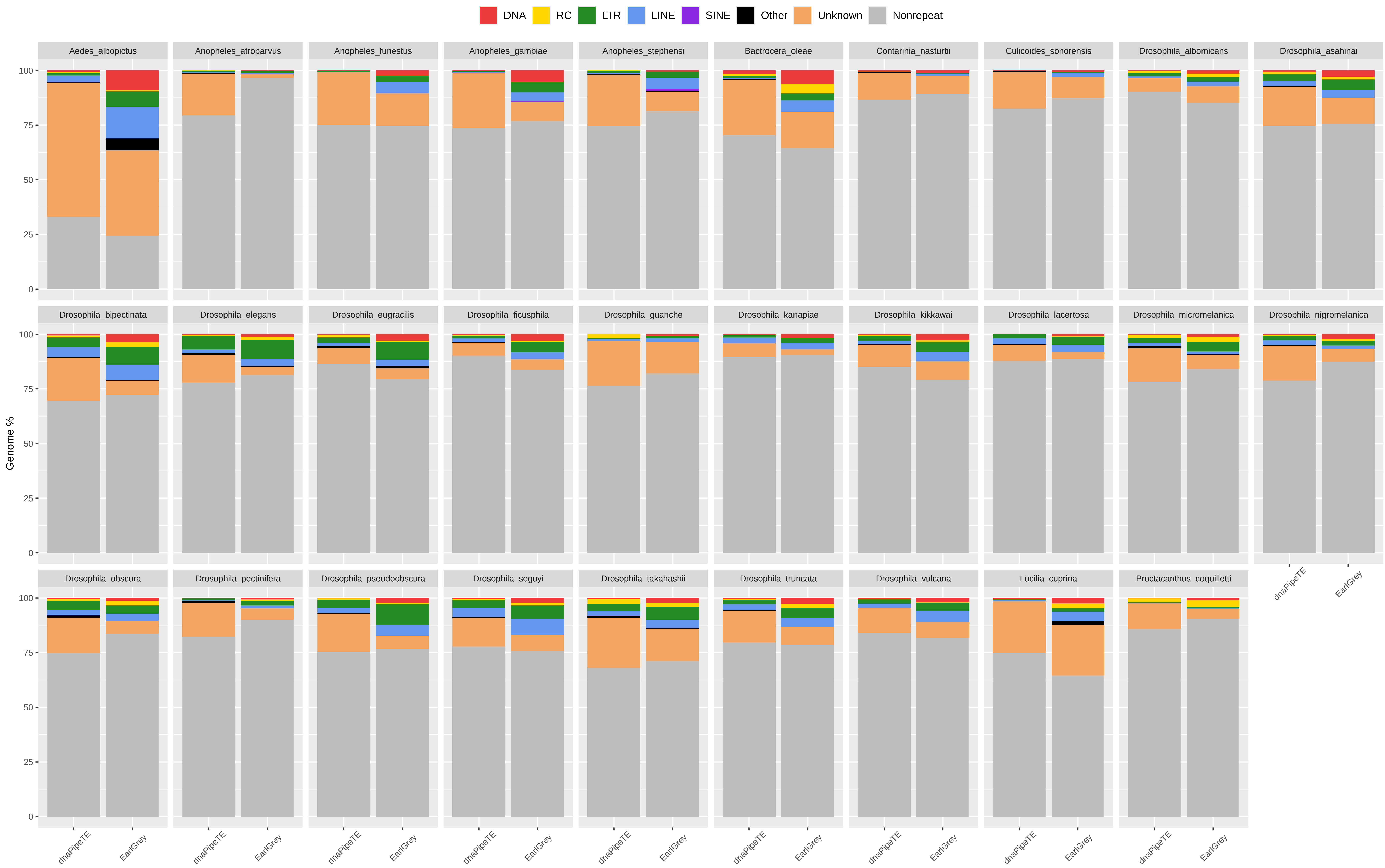


Supplemental Figure S2. Genomic proportion occupied by repeats in 29 dipteran species as estimated by EarlGrey and by the dnaPipeTE wrapper pipelines. The genome percentage is calculated proportionally to the assembly size in the case of EarlGrey, while it is calculated in relation to the genome size estimated in this study in the case of dnaPipeTE. DNA = DNA elements; RC = Rolling Circle; LTR = Long Terminal Repeats; LINE = Long Interspersed Nuclear Elements; SINE = Short Interspersed Nuclear Elements. 'Other' includes simple repeats, microsatellites, RNAs. 'Unknown' includes all repeats that could not be classified.
