## Supplemental Figure S3 for "Effective population size does not explain long-term variation in genome size and transposable element content in animals"

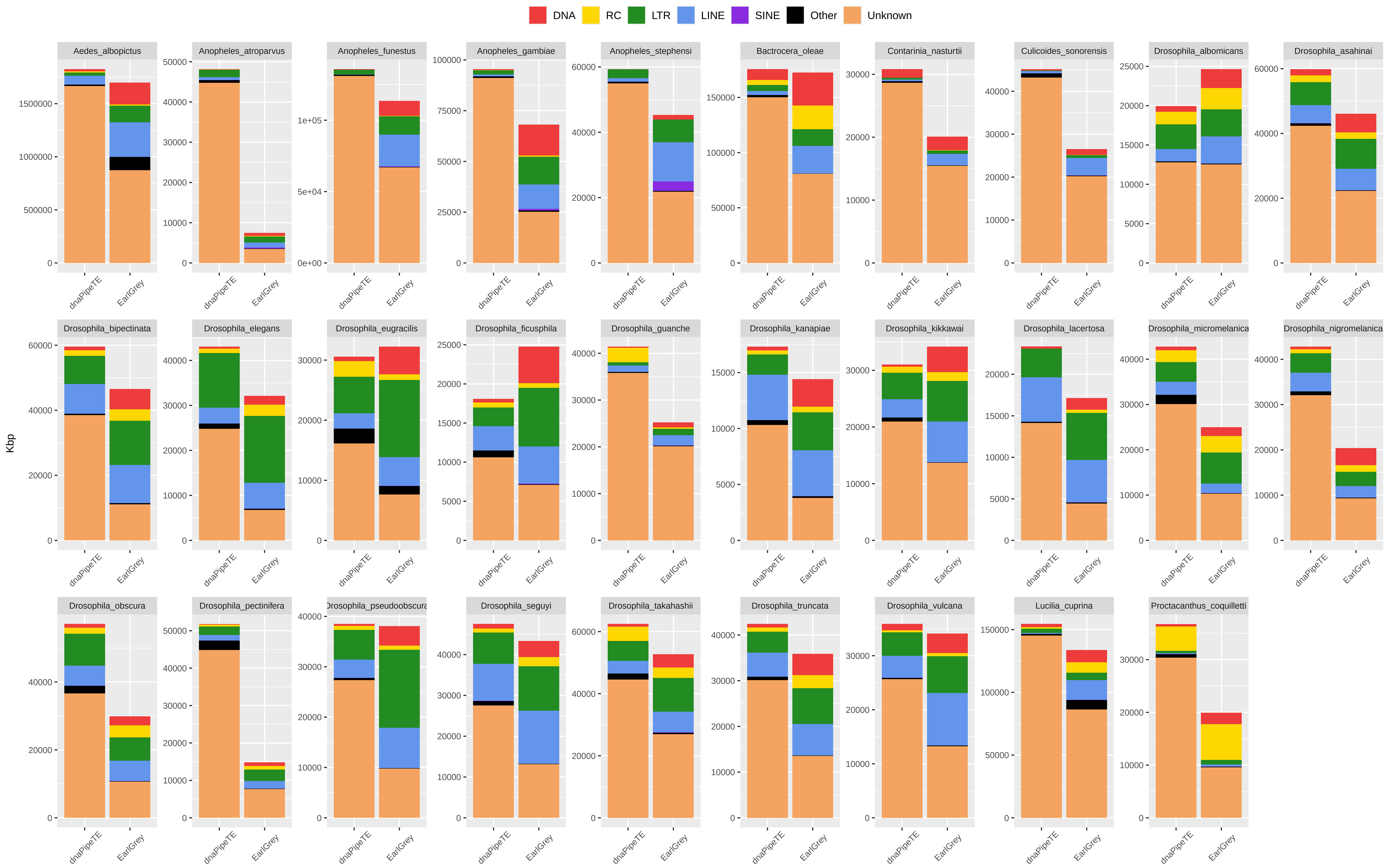


Supplemental Figure S3. Genomic proportion occupied by repeats in 29 dipteran species as estimated by EarlGrey and by the dnaPipeTE wrapper pipelines. Kbps are reported as the total base pairs annotated in the assembly in the case of EarlGrey, and as the estimated coverage from reads sampling in the case of dnaPipeTE. DNA = DNA elements; RC = Rolling Circle; LTR = Long Terminal Repeats; LINE = Long Interspersed Nuclear Elements; SINE = Short Interspersed Nuclear Elements. 'Other' includes simple repeats, microsatellites, RNAs. 'Unknown' includes all repeats that could not be classified.
