## Supplemental Figure S4 for "Effective population size does not explain long-term variation in genome size and transposable element content in animals"

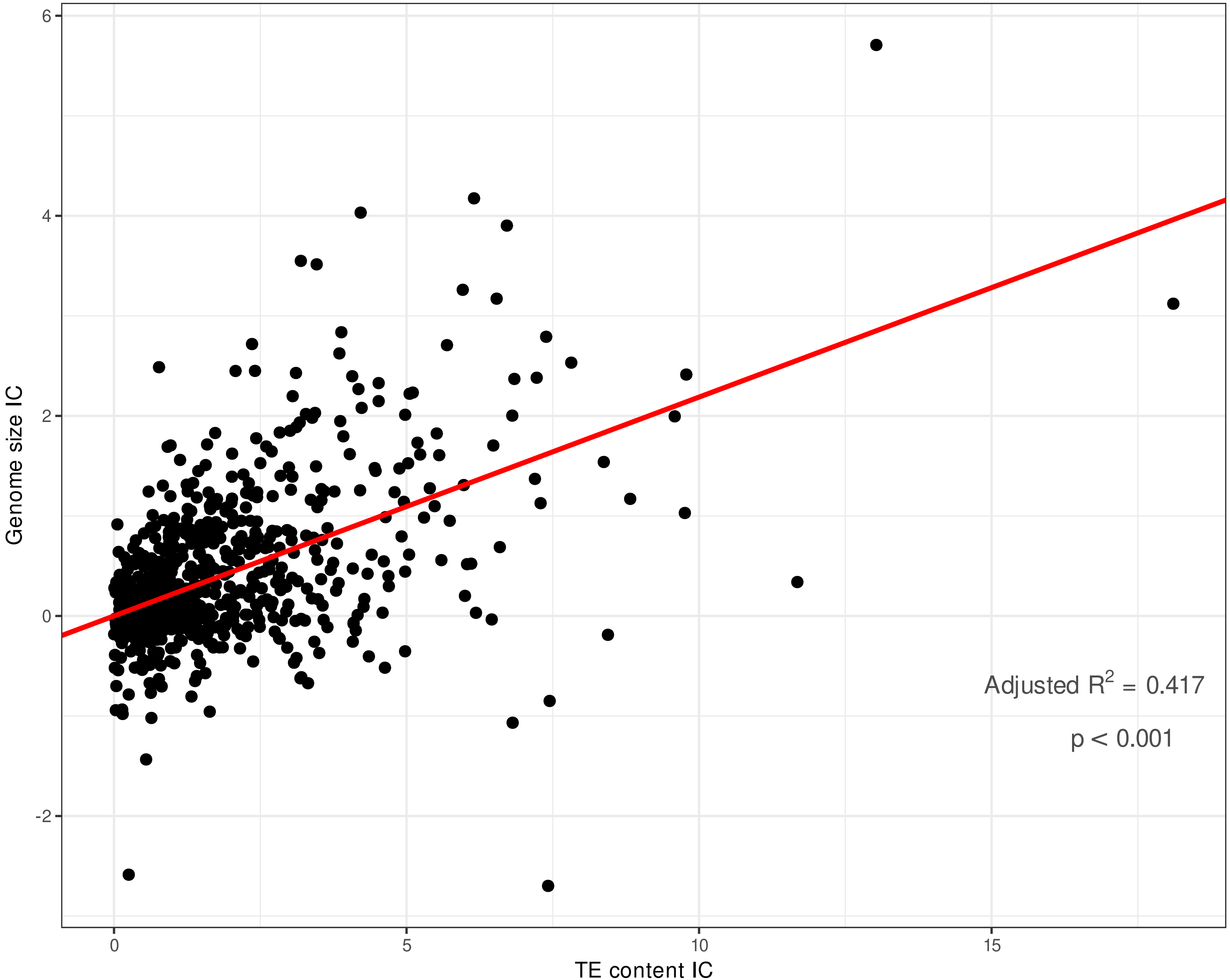


Supplemental Figure S3. Scatterplot of PIC regression for overall TE content as a predictor of genome size across the full dataset. Slope = 0.219, adjusted-R2 = 0.417, p-value < 0.001. Variables were log-transformed previous to regression.
