## Supplemental Figure S5 for "Effective population size does not explain long-term variation in genome size and transposable element content in animals"

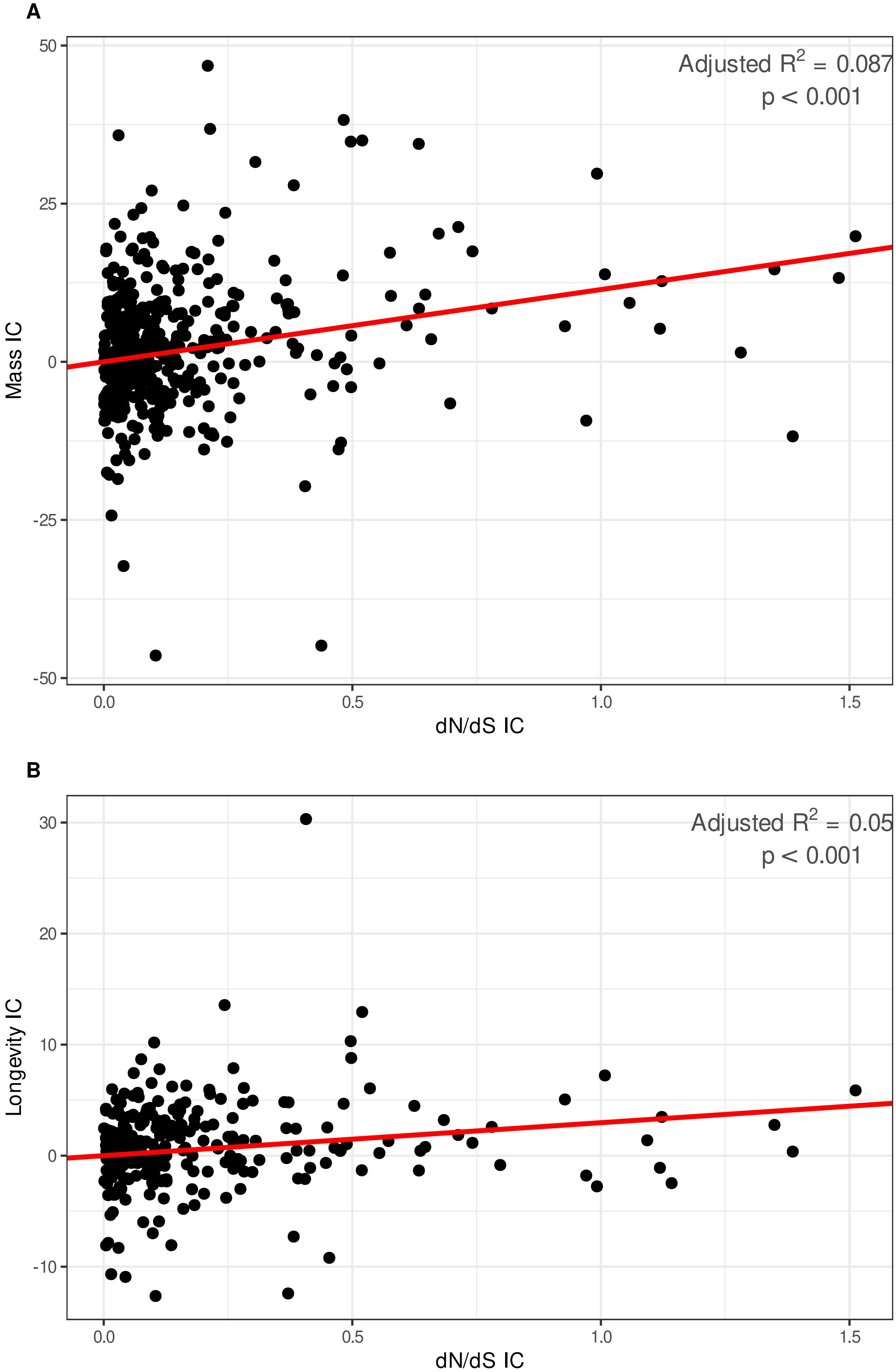


Supplemental Figure S4. Scatterplots of PIC regressions for Coevol dN/dS estimated from the GC3-poor geneset as predictor of (A) body mass and (B) longevity. Body mass: slope = 11.422, adjusted-R2 = 0.087, p-value < 0.001. (B) Longevity: slope = 2.970, adjusted-R2 = 0.050, p-value < 0.001. LHTs were log-transformed previous to regression.
