## Supplemental Figure S6 for "Effective population size does not explain long-term variation in genome size and transposable element content in animals"

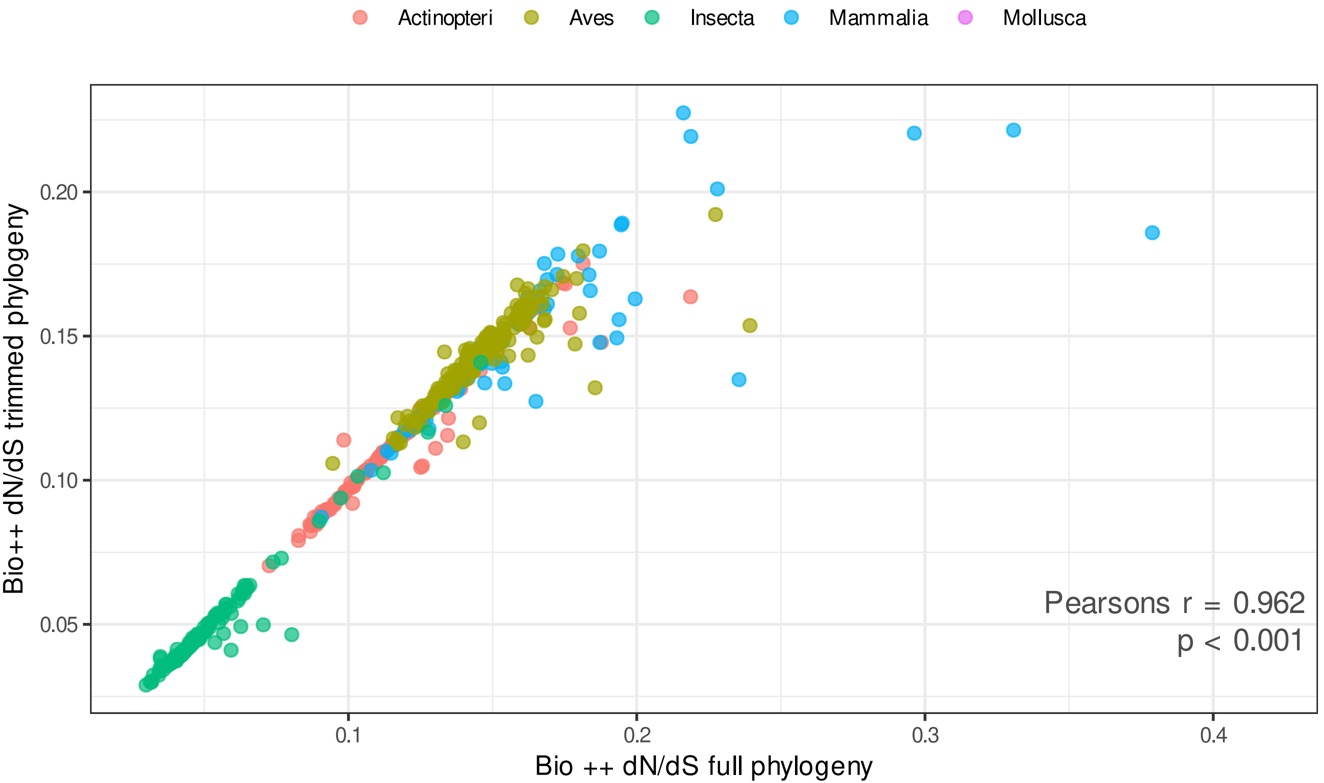


Supplemental Figure S5. Comparison of Bio++ dN/dS estimated from the full phylogeny and from the phylogeny where branches longer than 1 and shorter than 0.01 were pruned. Pearson's r = 0.962, p-value < 0.001.
