## Supplemental Figure S7 for "Effective population size does not explain long-term variation in genome size and transposable element content in animals"

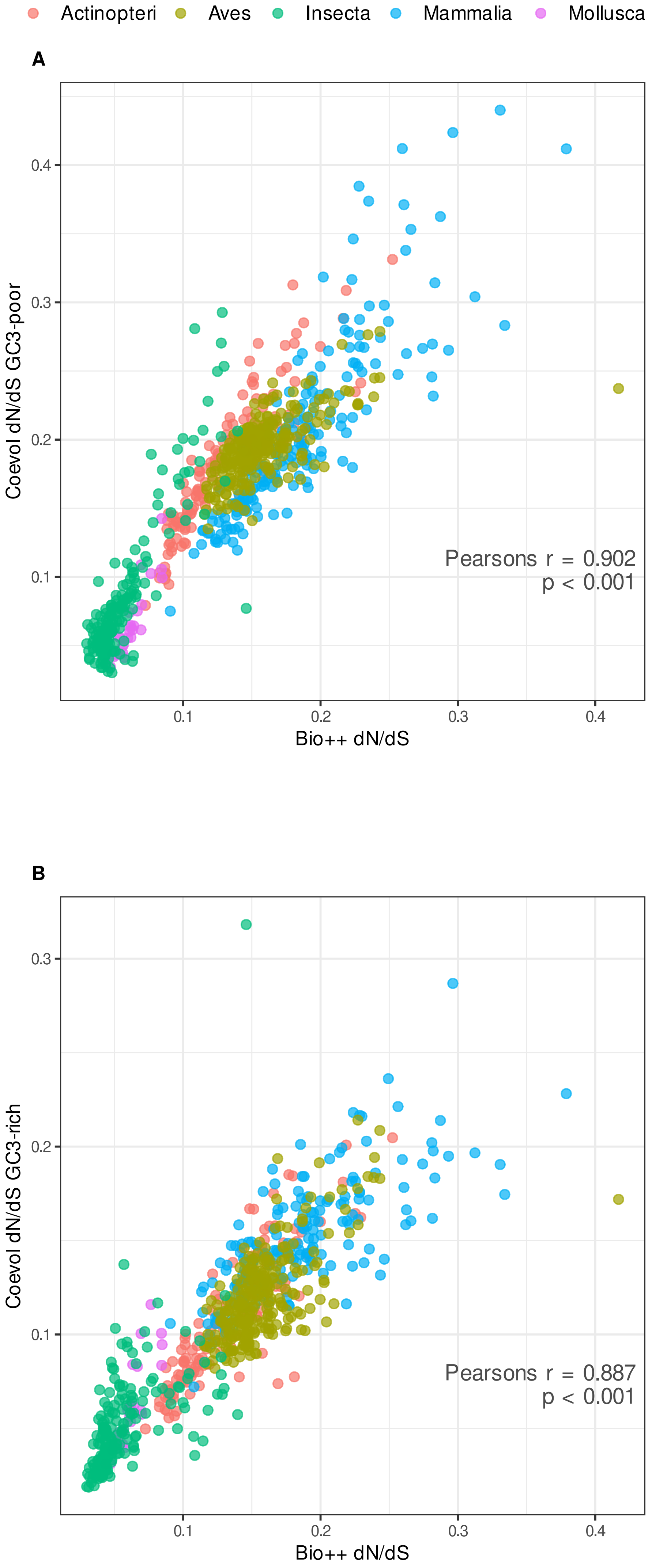


Supplemental Figure S6. Comparison of Bio++ and Coevol dN/dS estimations. Correlation coefficients are 0.902 and 0.887 using Coevol estimations over the GC3-poor (A) and GC3-rich (B) genesets, respectively (Pearson's r, p-value < 0.001).
