## Supplemental Figure S8 for "Effective population size does not explain long-term variation in genome size and transposable element content in animals"

A

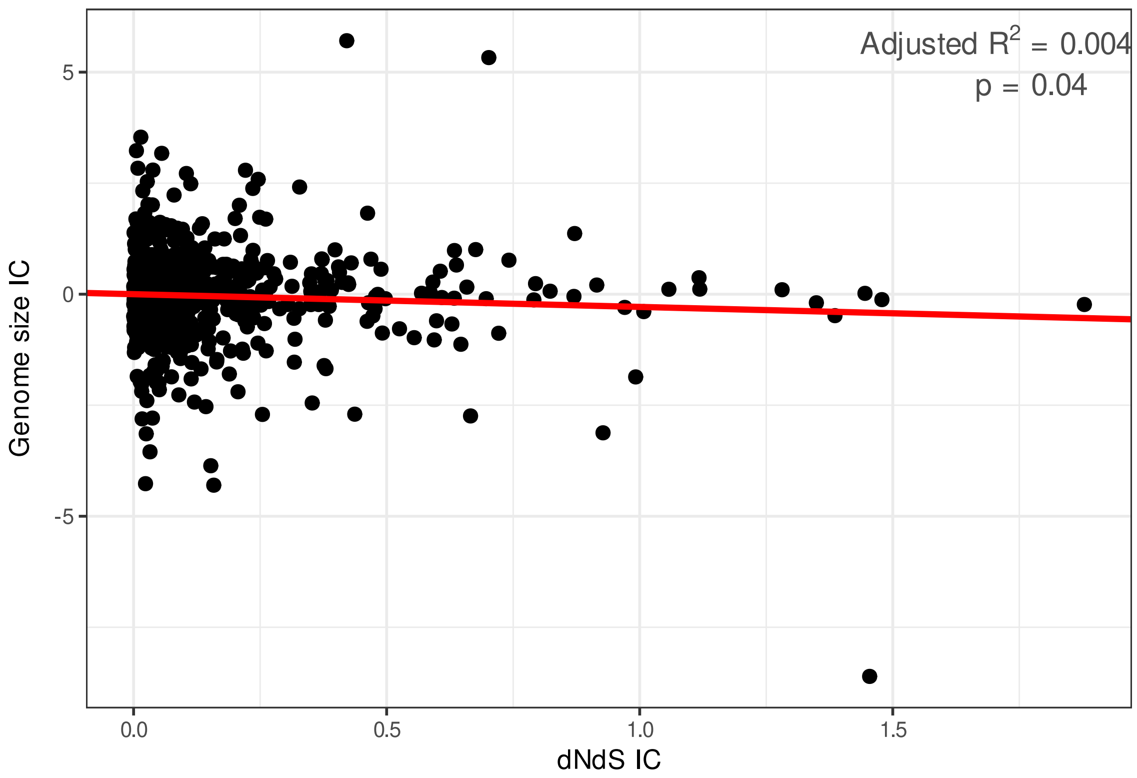


B

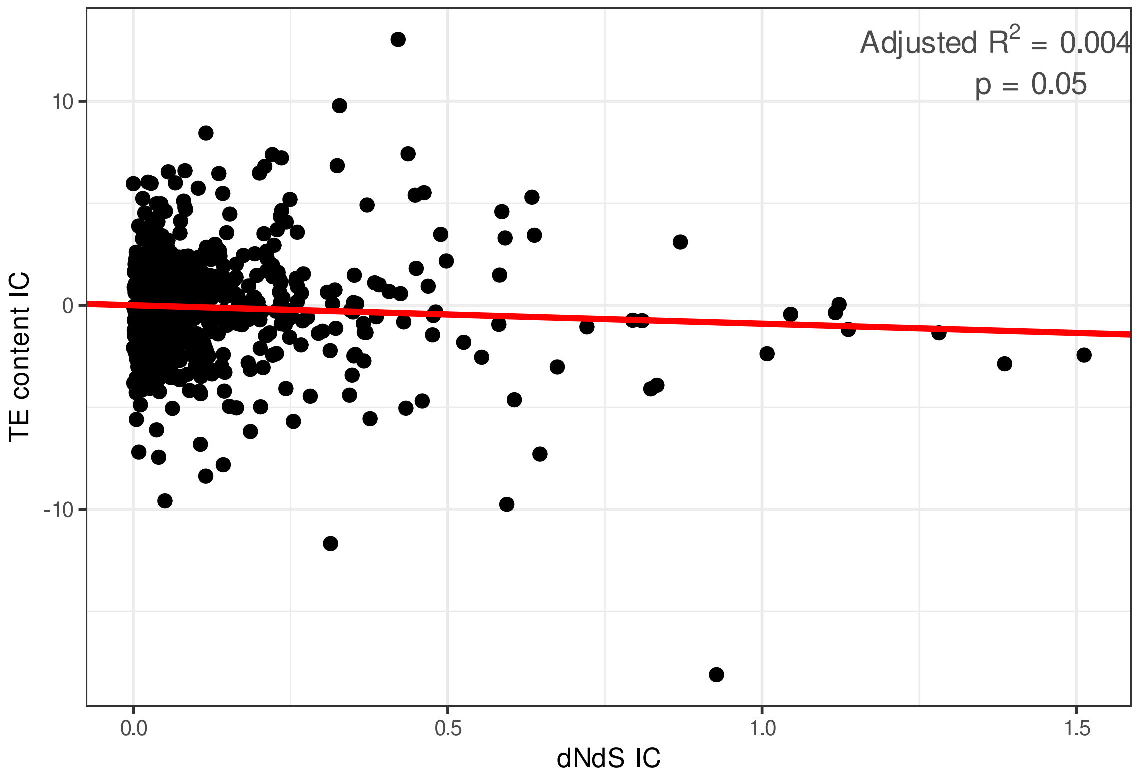


C

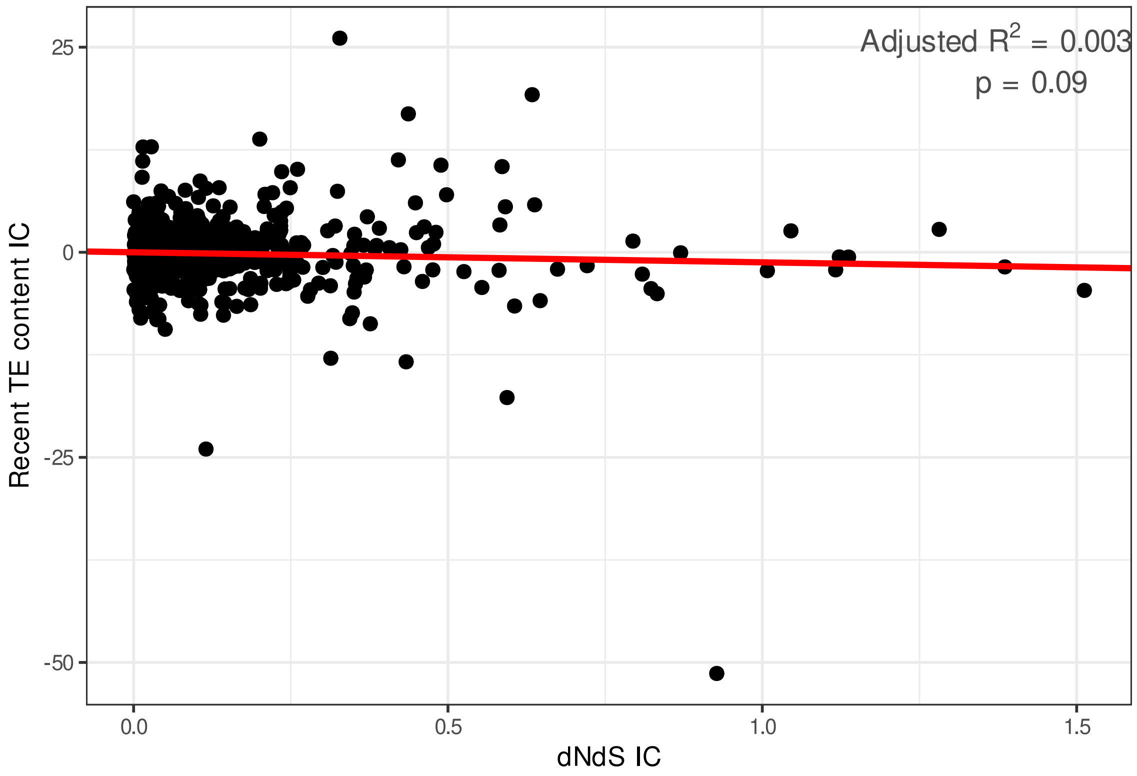


Supplemental Figure S7. Scatterplots of PIC regressions for Coevol dN/dS estimated from the GC3-poor geneset as predictor of (A) genome size, (B) overall and (C) recent TE content. Genome size: slope = -0.287, adjusted-R2 = 0.004, p-value = 0.039. Overall TE content: slope = -0.903, adjusted-R2 = 0.004, p-value = 0.050. Recent TE content: slope = -1.225, adjusted-R2 = 0.003, p-value = 0.089. Genomic traits were log-transformed previous to regression.
